# Cutaneous suction-mediated transfection in mice for delivery of DNA-encoded vaccines and proteins

**DOI:** 10.1101/2025.09.10.675275

**Authors:** Emran O. Lallow, Isabel Brandtjen, Yaxin Mo, Melissa Gulley, Louis Osorio, Sagar Kudchodkar, Nandita C. Jhumur, Christine C. Roberts, Lisa K. Denzin, David I. Shreiber, Biju Parekkadan, Hao Lin, Joel N. Maslow

## Abstract

An important step to fulfill the functionalities of DNA vaccines and therapeutics is transfection *in vivo* to produce the encoded antigens or therapeutic proteins. A cutaneous suction-based method has demonstrated effectiveness in many animal models and has been successfully applied in human clinical trials, but has not been extended to mouse models, where numerous disease models, transgenic strains, and murine-specific reagents exist. The current work establishes and optimizes methods for cutaneous suction-mediated DNA transfection in mice. By adapting a smaller cup diameter and smaller injection volume, the challenges of skin hyperelasticity and decreased skin thickness can be effectively addressed, and vaccinating mice with the GLS-5310 SARS-CoV-2 DNA vaccine yielded high levels of binding antibody and T cell responses. Additionally, suction following injection of a novel pVAX1-based expression vector yielded systemic levels of a SEAP transgene. Thus, suction-mediated delivery of nucleic acid-based therapies and vaccines can be a valuable tool for the study in pre-clinical mouse models.

## Introduction

DNA vaccines and therapeutics have demonstrated great promise due to their stability, efficacy, and cost-effectiveness [1-3], with one vaccine, ZyCoV-D, approved for human use in India [4]. Although several devices have been used to promote *in vivo* transfection of injected DNA, the currently available delivery mechanisms are associated with high-cost, tissue damage, and rigorous training requirements [1, 5, 6]. We developed a novel suction-based delivery device, GeneDerm, that has shown applicability in pre-clinical animal models, including hamsters, rats, rabbits, and ferrets, and has been successfully applied in human clinical trials of the GLS-5310 DNA vaccine [7-11]. The GeneDerm device is used to apply negative pressure suction atop the site of intradermally injected plasmid DNA, which in turn induces efficient *in vivo* DNA transfection. GeneDerm has many advantages as a DNA delivery device, such as being cost effective, pain-free, and requiring minimal user training. To date, GeneDerm has not yet been widely used in mice primarily due to the hyperelasticity of mouse skin and decreased skin thickness as compared to other pre-clinical animal models. As numerous transgenic strains and murine-specific reagents exist that are central to the pre-clinical study of many diseases, extension of this DNA delivery method to mice is highly desirable.

Here, we establish and optimize methods for suction-mediated DNA transfection in mice. We show that by using a smaller cup diameter and smaller injection volume, we are able to achieve high levels of transfection and transgene expression for DNA vaccines. Additionally, we show that the use of GeneDerm following the injection of a novel pVAX1-based expression vector yields detectable quantities of expressed proteins in the bloodstream of mice. GeneDerm can be a valuable tool for the study of nucleic acid-based therapies and vaccines in pre-clinical mouse models.

## Materials and methods

### Animal preparation and experiments

Six- to eight-week old BALB/c (BALB/cAnNTac) and C57BL/6 (C57BL/6NTac) female mice were purchased from Taconic Biosciences Inc. (Germantown, NY). Mice were housed under controlled conditions in accordance with the guidelines established by the Rutgers University Institutional Animal Care and Use Committee, under protocol IACUC-201800077.

Preparation of mouse skin for intradermal (ID) injection and subsequent application of suction was performed as previously described for rats and hamsters [10] with the following modifications (Figure 1). Following hair removal from the dorsal region by shaving and depilation (Figure 1A), the skin was pinched with the thumb and forefinger. Intradermal (ID) injection was accomplished with a 31-gauge SOL-M needle (SOL-Millennium, Chicago, IL) bevel upwards, parallel to the skin surface, ensuring very shallow injection (Figure 1B) to yield a post-injection bleb (Figure 1C) within the mouse skin. Suction is then applied atop the bleb using the GeneDerm device with a cup opening of either 2 mm or 3 mm (Figure 1D), as noted.

**Figure 1.**
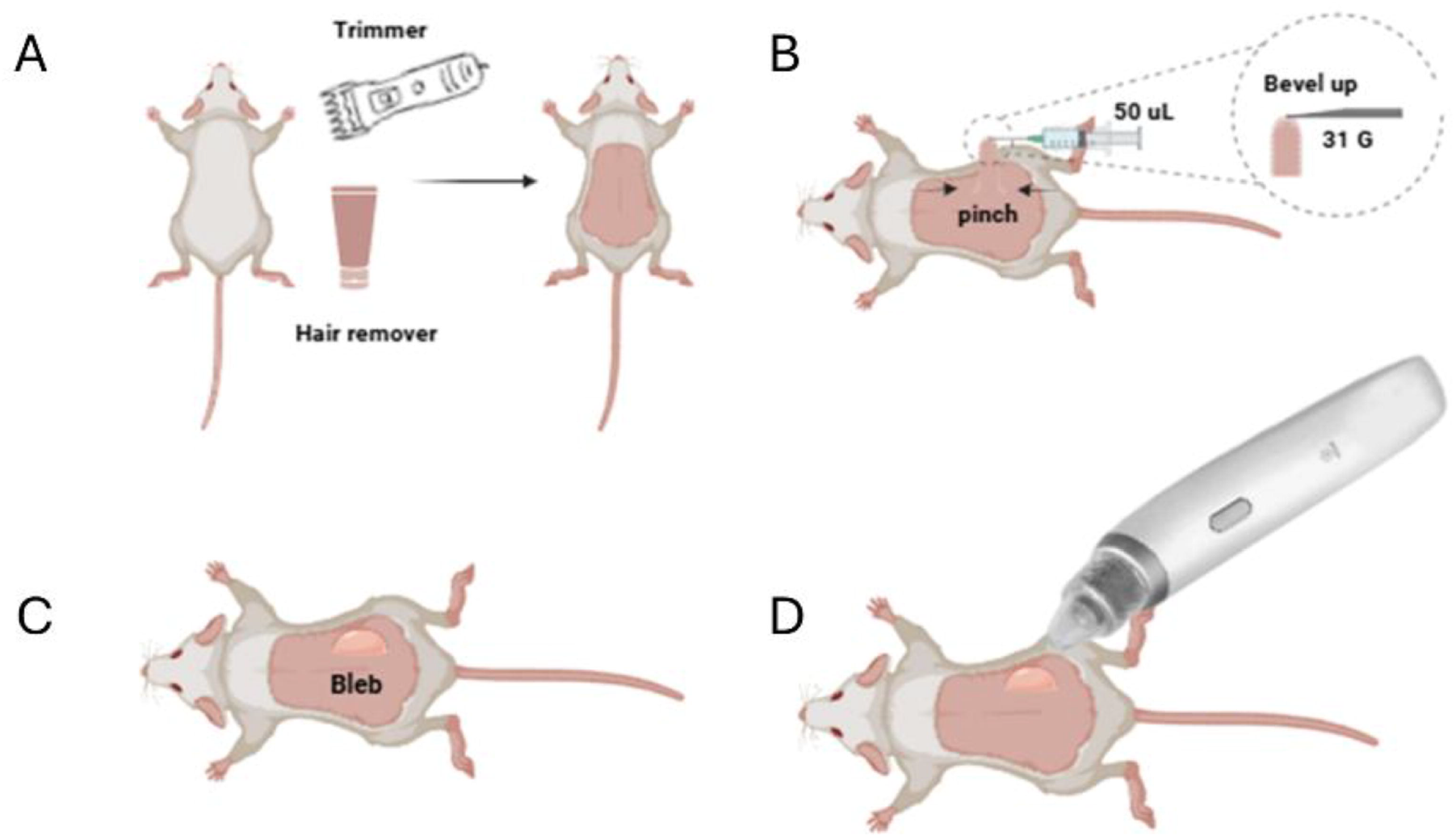
Schematic diagram of the experimental procedure and technique. (A) The hair is removed from the dorsal area. (B) A solution is injected intradermally into pinched skin to form the bleb. (C), (D) The application of the negative pressure using the GeneDerm Device atop the bleb.

### Luminescence imaging and quantification

Six- to eight-week old BALB/c (BALB/cAnNTac) and C57BL/6 (C57BL/6NTac) female mice were injected with pGLS-Luc2 (purchased as pGL4.14, Promega, Madison, WI), a plasmid encoding the firefly luciferase protein. XenoLight D-Luciferin Potassium Salt (PerkinElmer, Waltham, MA) was injected intraperitoneally (IP) at a dose of 150 mg/kg in a 200 µL volume. At 10 minutes post-injection, animals were placed under anesthesia and imaged using the IVIS Lumina X5 live imaging system (PerkinElmer, Waltham, MA). Luminescence was determined following a 3-second exposure. Aura imaging software (Model 4.0.8, Spectral Instruments, Tucson, AZ) was used to quantify the total photon emission at each injection site. Background signal was determined for a non-treated region and then subtracted from the treated regions. A square root transformation was applied to all conditions. Graphical presentation and statistical analysis were performed using GraphPad Prism (10.2.3, GraphPad Software, San Diego, CA).

### GLS-5310 vaccination and immunology

Ten six- to eight-week old BALB/c (BALB/cAnNTac) mice were vaccinated with 25 µg of the GLS-5310 COVID-19 DNA vaccine, which expresses the SARS-CoV-2 spike and ORF3a proteins [7], in a 50 µL volume on days 0 and 14. Immediately following ID injection, suction was applied to the injection sites using the GeneDerm device (80 kPa, 30 sec) using a 3-mm cup opening. Blood was collected from the retro-orbital plexus prior to vaccination and via cardiac puncture on day 28. Sera was stored at -20°C for enzyme linked immunosorbant assay (ELISA) analysis against the SARS-CoV-2 spike protein as described [7]. Spleens were collected on day 28 and immediately processed to harvest splenocytes for enzyme-linked immunospot (ELISpot) analysis as described [7].

### In vitro transfection of HaCaT cells

A custom pVAX1-based vector was designed using VectorBuilder (Chicago, IL) to include a strong cytomegalovirus (CMV) promoter that constitutively activates secreted embryonic alkaline phosphatase (SEAP) and was designated as pV1-SEAP. To verify the performance of the new expression vector *in vitro*, adult human skin keratinocyte cells (HaCaT) cells (gift from the lab of Francois Berthiaume at Rutgers University) were maintained using DMEM/F12 with 10% FBS and 1% anti-anti (ThermoFisher, Waltham MA). HaCaT cells were transfected using FuGENE 6 (Madison, WI) and purified pV1-SEAP DNA at a 3:1 ratio of FuGENE to DNA. Plasmid DNA (0.5 µg, 1 µg, 2 µg, 4 µg, and 6 µg) was incubated for 30 minutes at room temperature with 6 µL of FuGENE 6 and Opti-MEM media (ThermoFisher, Waltham MA) to create DNA complexes. These complexes were then added dropwise to a 6-well plate containing 250,000 HaCaT cells in DMEM/F12 and then incubated at 37 °C in 5% CO2 overnight. The media was changed to fresh DMEM/F12 24-hours post transfection, and supernatants were collected at baseline, after 24 hours, and after 48 hours, and stored at -20 °C. For SEAP determination, samples were thawed on ice, and SEAP was quantified using a Phospha-Light SEAP Reporter Gene Assay System and plate reader (VarioSkan LUX, ThermoFisher, Waltham MA). Results were plotted and analyzed using GraphPad Prism.

### Perfusion of transfected HaCaT cells

Three wells of a 6-well plate containing HaCaT cells (250K cells) were transfected with pV1-SEAP as above, with non-transfected wells serving as a control. Cells were perfused with DMEM/F12 media for 60 hours at a flow rate of 0.5 mL/hour into the well with fractionated supernatant collected every two hours, as previously described [12]. SEAP concentrations were determined using the Phospha-Light SEAP Reporter Gene Assay System (ThermoFisher, Waltham MA) and results analyzed for dynamic transgene activation dynamics using GraphPad Prism.

### In vivo transfection with pV1-SEAP and analysis

On week 1, Balb/c mice were sampled for a baseline reading of plasma SEAP levels by sampling 100 µL of whole blood. The plasma was separated then stored in -20 °C. At week 2, the mice were injected ID with 20 µg pV1-SEAP in a 50 µL injection volume in four regions in the dorsal area. Immediately following injection, suction was applied to each injection site using the GeneDerm device as above. At 24- and 72-hours post-injection, whole blood was collected, and plasma was separated to quantify circulating SEAP levels. The data was normalized to the background and to the total volume of the sample. Data was plotted and analyzed using GraphPad Prism.

## Results

### Optimization of suction-mediated in vivo transfection in mice with long-term transgene expression

Our prior studies showed that suction-mediated *in vivo* transfection of plasmid DNA in rat skin was dependent on induced tissue strain and tension, which in turn depended on the applied suction pressure and opening diameter of the GeneDerm cup. Transfection was largely independent of the time that suction was applied [10, 13]. The marked laxity of mouse skin prevented successful extension of the current suction protocol to mice (data not shown), requiring modification of the methodology for injection and application of suction. To assess and optimize transgene expression of DNA plasmids following ID injection with suction mediated transfection in mice, expression from a luciferase-encoded plasmid was studied.

To optimize the conditions for GeneDerm use in mice, we assessed 4 separate parameters. The first was to optimize the method for mouse ID injections. Using a Mantoux injection technique, as we had done for other animals and humans, we found that creating a taut fold of skin through a gentle pinch and lift and then injecting into the superior aspect of skin along the raised fold yielded a fluid-filled bleb akin to that observed in our other studies [10, 14] and indicating successful ID injection (Figure 1).

We next assessed the effects of the cup opening on the GeneDerm device through which suction is delivered to the skin. In early attempts to apply GeneDerm in mice, the 6-mm diameter opening used in all other species [7-11] was too large for use in mice, as their comparatively looser skin was pulled into the cup and yielded no detectable luminescence (data not shown). Therefore, we used a cup opening diameter of either 2 or 3 mm. We assessed luciferase expression in Balb/c mice following injection of pGLS-Luc2 either alone (as control) or followed by application of 80 kPa of suction for 30 sec. Luminescence 24-hours post-injection was greater for conditions with applied suction, but it was only significantly different from injection-only control when the 3-mm cup opening was employed (Figure 2). A qualitative assessment of the time course of luciferase expression using a 3-mm cup opening (Figures S1) showed that luminescence was detectable as early as 1-hour post-injection, was the highest from 1 through 9 days, and remained detectable up to 22 days post-injection. At each time point, pGLS-Luc2 administered with suction yielded higher levels of luminescence than injection only. A similar preliminary assessment of suction-mediated transfection in C57bl/6 mice was only minimally successful (data not shown) as luminescence detection is significantly decreased in this species due to their dark fur and skin, as reported by others [15].

**Figure 2.**
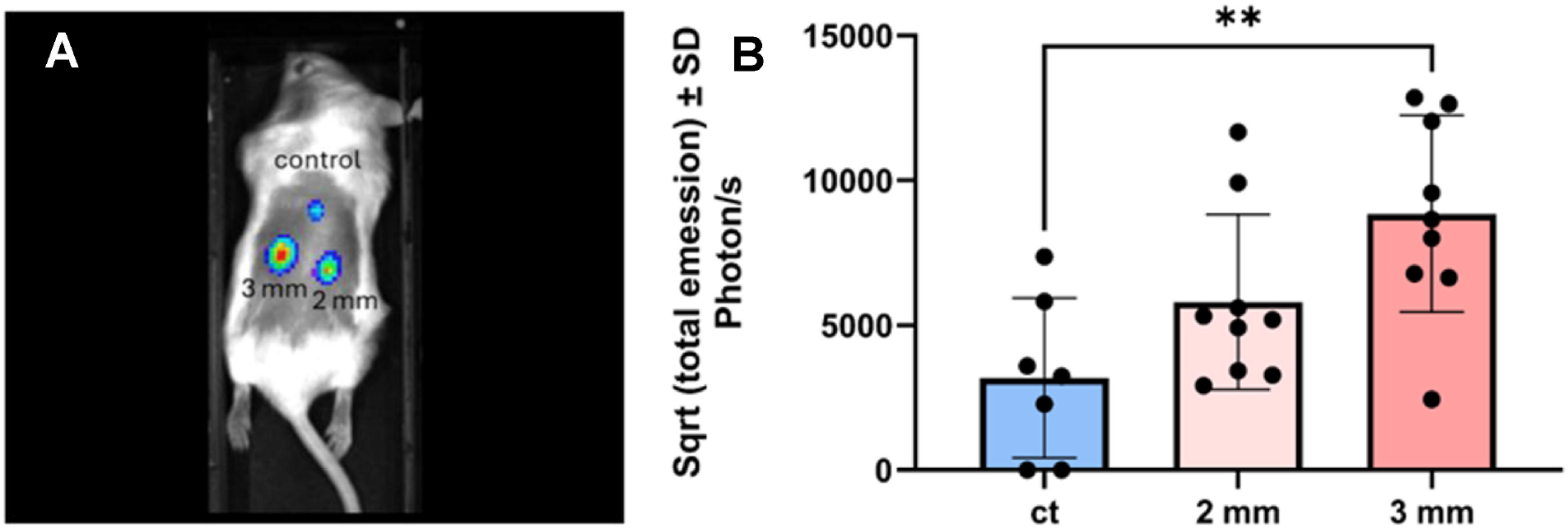
Luminescence expression in Balb/c mice using different cup diameters at various imaging time points. 50 µL of pGLS-Luc2 was injected intradermally for all animals with or without suction pressure application; suction pressure was 80 kPa applied for 30 seconds using the GeneDerm device. (A) Representative luminescence image with 3 seconds exposure time. (B) Luminescence quantification comparing 2 and 3-mm cup diameters to injection only control 1-day post treatment. Data represents mean±SD. Statistics are presented as **p ≤ 0.01 by one-way ANOVA followed by a Tukey’s multiple comparisons test.

Finally, we assessed whether total injection volume or the strength and time of suction affected luciferase expression. Balb/c mice were injected with 25 µg of pGLS-Luc2 as either a 10-µL volume at a concentration of 2.5 µg/µL, a 30-µl volume at a concentration of 0.83 µg/µl, or a 50-µL at a concentration of 0.5 µg/µL. When suction was applied, there were no differences among these three conditions (Figure 3A). Finally, we found no significant difference in transgene expression when applying 65 kPa of suction for 15 seconds, 80 kPa for 15 seconds, or 80 kPa for 30 seconds with the GeneDerm device (Figure 3B).

**Figure 3.**
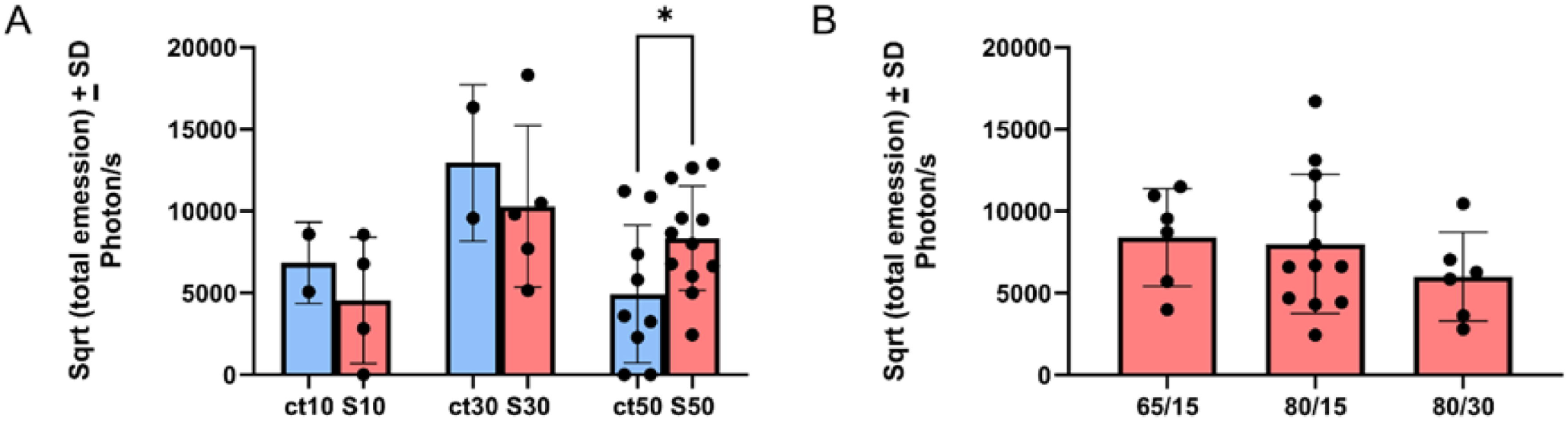
Effects of varying injection volumes, pressure and time using Balb/C mice. (A) A comparison of injection only control, “ct”, at volumes of 10, 30, and 50 µL with suction pressure applied at 80 kPa for 30 seconds. (B) A 50-µL injection was used to compare two pressures, 65 kPa and 80 kPa, applied for 15 or 30 seconds. Data represents mean±SD of 3 seconds exposure time of total luminescence emission. Statistics are presented as *p ≤ 0.05 by one-way ANOVA followed by a Tukey’s multiple comparisons test.

### Immune responses for mice following suction-mediated transfection with GLS-5310 COVID DNA vaccine

As a further test of suction-mediated *in vivo* transfection of mice for vaccination, we determined the immune response of Balb/c mice vaccinated with the GLS-5310 SARS-CoV-2 DNA vaccine. Binding antibody titers against the SARS-CoV-2 spike protein for mice vaccinated with suction were approximately 1.5 log greater 14 days post second vaccination than injection-only controls (p<0.001; Figure 4A). Additionally, all 10 suction-treated animals seroconverted, versus 8 (80%) of injection-only animals, two of which had only borderline responses. Vaccination with GLS-5310 and GeneDerm suction demonstrated a broad-based T-cell response against both the spike and ORF3a proteins (Figure 4B) that was more than 2-fold higher (213.6 versus 90.7 for the total SFU summing up all pools; p ≤0.05) for animals vaccinated without suction.

**Figure 4.**
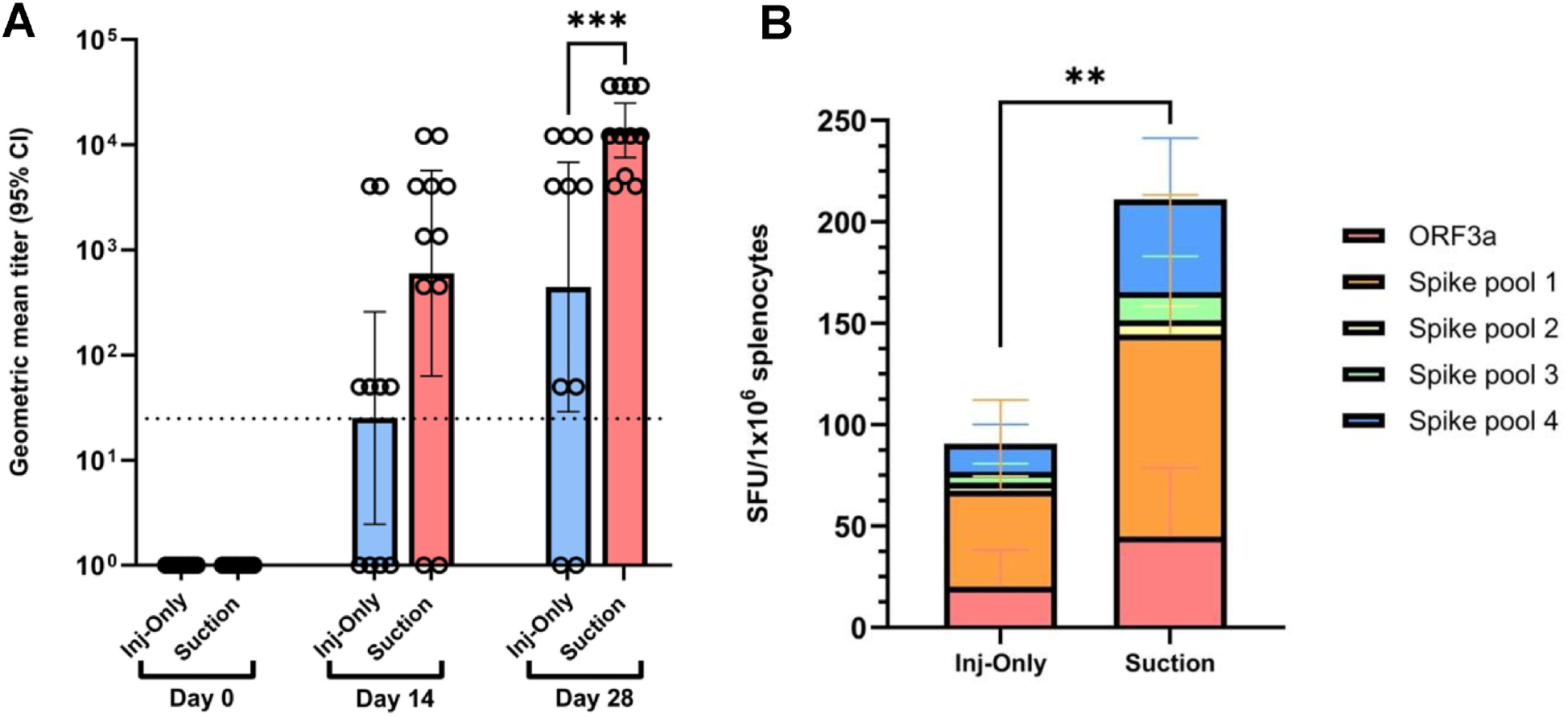
Immune responses to a SARS-CoV-2 DNA vaccine candidate using Balb/c mice. (A) Antibody responses of 50 µL injections of injection only control compared to 80 kPa pressure applied for 30 seconds with a 3-mm diameter cup. (B) T-cell responses quantified via spot forming units (SFU) by an ELISPOT assay to pools of peptides spanning the Spike and ORF3a proteins. Statistics are presented as **p ≤ 0.01 and ***p ≤ 0.001 by ordinary one-way ANOVA followed by a Tukey’s multiple comparisons test. Statistics in (B) are based on the total number of SFU by summing up contributions from the pools.

### In vitro and in vivo measurement of SEAP expression

Initial experiments were performed to optimize *in vitro* transfection of pV1-SEAP and subsequent expression of SEAP in HaCaT cells by varying the amount of purified plasmid pV1-SEAP DNA (Figure S2). The most efficient condition was using 2 µg of pV1-SEAP plasmid DNA, which yielded a four-log increase in SEAP production relative to control cells (Figure 5A and supplementary data). We then determined the time course of SEAP expression in a kinetic assay perfusion system. Timescale dynamics showed a peak occurring at roughly 16 hours with high levels of expression until 30 hours followed by an exponential decay (Figure 5B).

**Figure 5.**
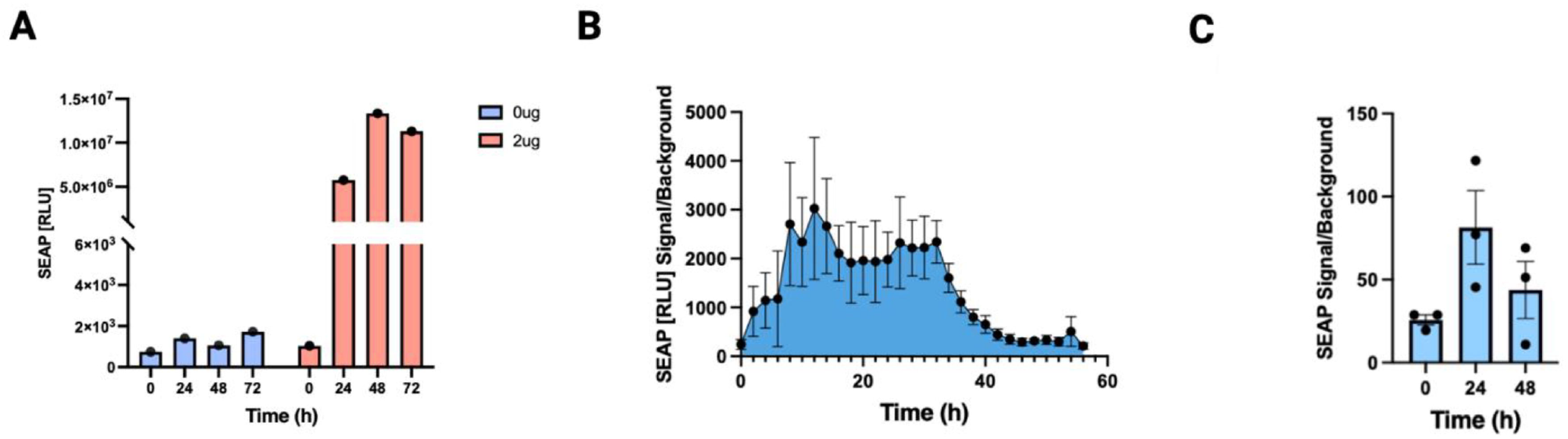
(A) In vitro transfection of HaCaT cells with 2 µg of pV1-SEAP DNA compared to a control produces 4-log increase in SEAP production. (B) 60-hour kinetic perfusion assay of transfected HaCaT cells using 2 µg of pV1-SEAP DNA illustrating the timescale dynamics of the transgene. (C) Balb/c mice injected with 25 µg pV1-SEAP DNA subcutaneously followed by 80 kPa of suction for 30 seconds with whole blood collection taken before the injection, 24 hours-post injection, and 48-hours post injection. The results were normalized to the background and volume of sample. A significant increase in SEAP production 24-hours post transfection is present.

We then investigated whether and to what extent mice transfected with pV1-SEAP would have detectable levels of expressed SEAP present in samples of circulating blood. Three Balb/c mice were injected with 25µg of purified pV1-SEAP DNA in a 50 µL volume, followed by application of suction using the GeneDerm device (80kPa, 30 sec) using a 3-mm cup size. Peak values were 3-fold greater than pre-treatment, with a peak at 24 hours, decreasing to approximately 1.5 times baseline at 48 hours (Figure 5C).

## Discussion

Suction-mediated *in vivo* transfection was originally developed in a rat model, which was deemed a more suitable model because the mechanical and physical properties of rat skin are more comparable to that of humans [16-19]. This similarity was evident, as the parameters used in rat skin were directly translated into use in human clinical trials without change [3, 7, 8]. However, mouse models are generally a more suitable model for early pre-clinical studies due to the availability of a wide range of supporting reagents, assays and techniques [20-23]. However, since the mechanical properties of mouse skin differ from rats and humans, adaptations of the method of ID injection, volume of material injected, and the diameter of the opening of the suction cup through which suction is transmitted were required. ID injection was successfully achieved using a smaller gauge needle (31 versus 28 gauge) and holding a fold of skin between the thumb and forefinger as demonstrated in Figure 1. Injection volumes were decreased to 50 µL from 100 µL, and we used a suction cup with an opening diameter of 3 mm instead of the 6 mm opening diameter used for other animals and humans. As shown for rats [10], pressure and time-of-suction had minimal effect on transgene expression.

To validate the results obtained after transfection with a vector encoding luciferase, we immunized mice with the GLS-5310 SARS-CoV-2 DNA vaccine. Suction significantly enhanced both the antibody and the cellular immune responses compared to the injection-only group, consistent with our prior studies in rats and hamsters [7, 10].

Finally, we showed that suction-mediated *in vivo* transfection of the skin yielded systemic levels of an expressed transgene. We first established that pV1-SEAP, a pVAX1 expression vector with a constitutive CMV promoter, transfected into human HaCaT skin cells *in vitro* yields high levels of SEAP expression. *In vivo* transfection of mice using suction showed transgene activation dynamics similar to the *in vitro* model. Considering the fact that the serum half-life of SEAP is approximately two hours [24], our data is consistent with constitutive production of SEAP in vivo.

## Conclusion

In conclusion, in this study we have established and optimized an intradermal suction-mediated DNA delivery method for mice. We additionally demonstrate that *in vivo* suction-mediated transfection of a COVID-19 DNA vaccine was highly immunogenic and that transfection of a SEAP expressing plasmid yielded high levels of circulating protein detected in blood with a pharmacodynamic profile similar to that seen *in vitro*.

## Author contributions

Emran O. Lallow, David I. Shreiber, Biju Parekkadan, Lisa Denzin, Hao Lin, and Joel N. Maslow conceived and designed the experiments for this study. Emran O. Lallow, Isabel Brandtjen, Yaxin Mo, Louis Osorio, Mellisa Gulley, Sagar Kudchodkar, and Nandita C. Jhumur performed the experiments. Hao Lin, Joel N. Maslow, Lisa K. Denzin, David I. Shreiber, and Biju Parekkadan provided resources and supervision. Emran O. Lallow and Isabel Brandtjen wrote the original concept and manuscript, and Hao Lin, Joel N. Maslow, Christine C. Roberts, Lisa K. Denzin, David I. Shreiber, Biju Parekkadan, and Yaxin Mo reviewed and edited it.

## Conflict of interest

Emran O. Lallow, Melissa Gulley, Sagar Kudchodkar, Christine C. Roberts and Joel N. Maslow are employees of GeneOne Life Science Inc.

The remaining authors declare that the research was conducted in the absence of any commercial or financial relationships that could be construed as a potential conflict of interest.

## Ethics statement

The animal study was reviewed and approved by Rutgers University Institutional Animal Care and Use Committee under protocol IACUC-201800077.

## Data availability statement

The data that support the findings of this study are available from the corresponding author upon reasonable request.

## Notes

### Competing Interest Statement

The authors have declared no competing interest.

