## Supplementary figures and images for "Cutaneous suction-mediated transfection in mice for delivery of DNA-encoded vaccines and proteins"

### Supplemental Figure S1

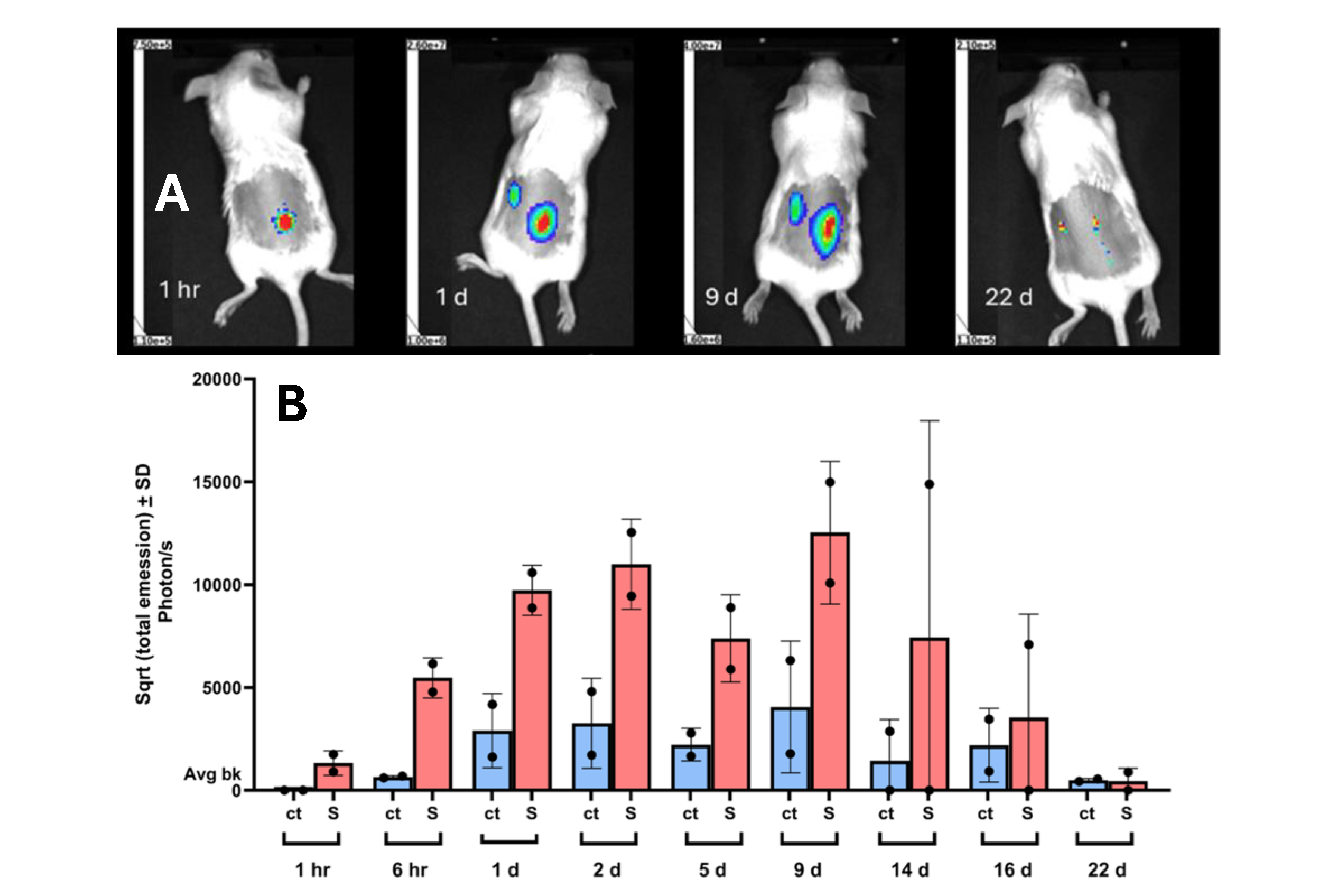

### Supplemental Figure S2

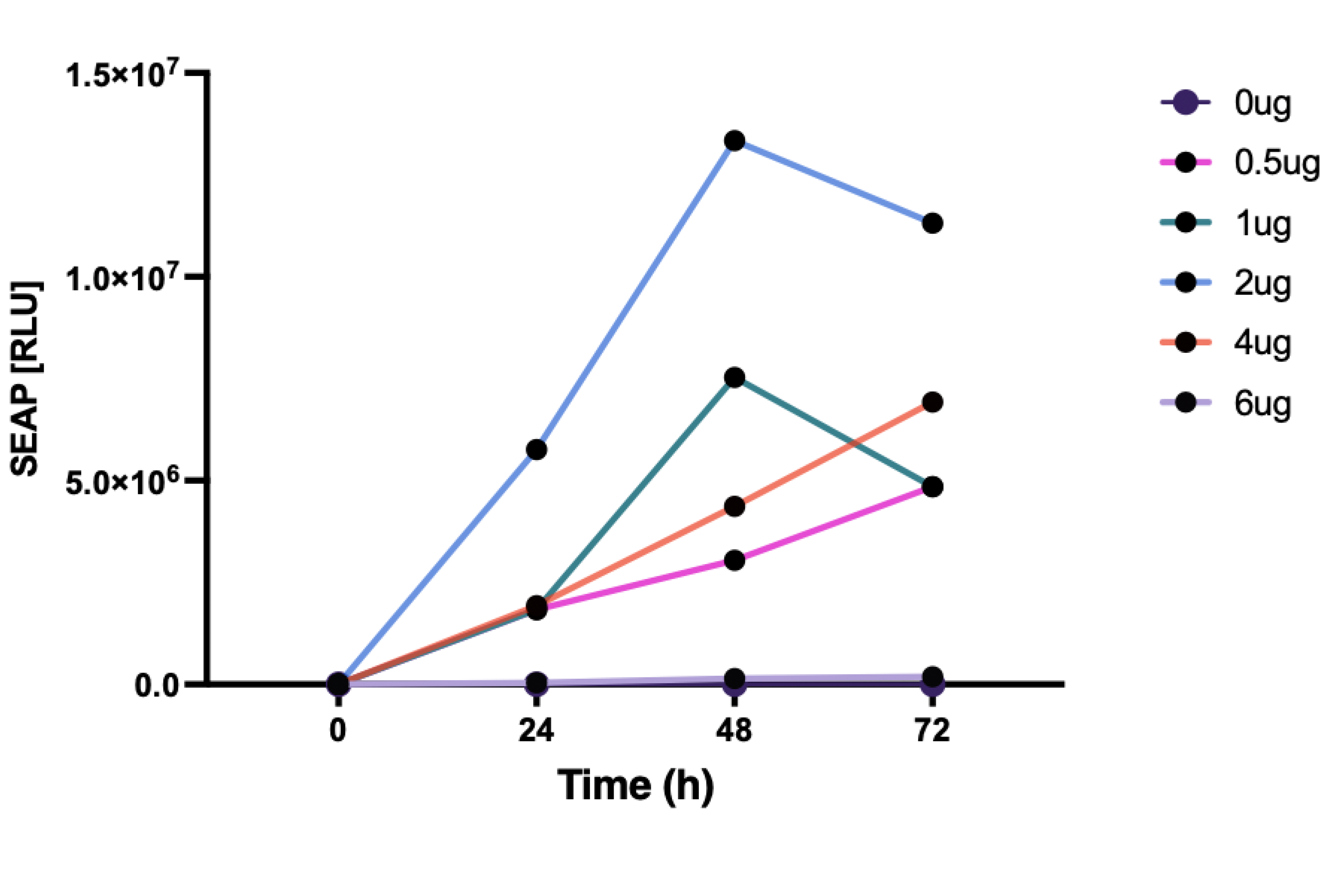
